# On the fitness effects and disease relevance of synonymous mutations

**DOI:** 10.1101/2022.08.22.504687

**Authors:** Xukang Shen, Siliang Song, Chuan Li, Jianzhi Zhang

## Abstract

We recently measured the fitness effects of a large number of coding mutations in yeast under four laboratory conditions, finding that most synonymous mutations are strongly deleterious although they are overall significantly less detrimental than nonsynonymous mutations. Kruglyak *et al*. believe that most nonsynonymous and nearly all synonymous mutations have no detectable fitness effects, so hypothesize that our results largely reflect the fitness effects of CRISPR/Cas9 off-target edits and secondary mutations that occurred in mutant construction. Dhindsa *et al*. argue that our findings contradict other yeast and human mutagenesis studies, human allele frequency distributions, and disease gene mapping results. We find Kruglyak *et al*.’s hypothesis unsupported by prior yeast genome editing studies and mutation rate estimates. Furthermore, their hypothesis makes a series of predictions that are falsified by our published and newly collected data. Hence, their hypothesis cannot explain our observations. Dhindsa *et al*.’s comparisons between synonymous and nonsynonymous mutations in prior mutagenesis studies and in contributions to disease are unfair and human allele frequency distributions can be compatible with our fitness estimates when multiple complicating factors are considered. While our fitness estimates of yeast synonymous mutants overturn the (nearly) neutral assumption of synonymous mutations, they are not inconsistent with various existing data.

## INTRODUCTION

We recently constructed 8,341 mutants each carrying a synonymous, nonsynonymous, or nonsense mutation in one of 21 representative genes of the yeast *Saccharomyces cerevisiae* [1]. We found that most synonymous and most nonsynonymous mutants are significantly less fit than the wild type, although nonsynonymous mutations are overall more detrimental than synonymous ones [1]. Kruglyak *et al*. believe that “most nonsynonymous and nearly all synonymous mutations have no or minimal effects on fitness” and “only a subset of mutations, primarily nonsynonymous ones, would have significant effects on fitness” [2]. They hypothesize that our findings largely reflect the fitness effects of CRISPR/Cas9 off-target edits and secondary mutations that occurred in mutant construction [2].

Our finding that most synonymous mutations are strongly deleterious (selection coefficient *s* < -0.001) led to the suggestion that human synonymous mutations may be substantially more important as a cause of disease than is currently thought [1]. Dhindsa *et al*. [3] argue that, compared with nonsynonymous mutations, human synonymous mutations are significantly less deleterious and are substantially less likely to cause disease. Furthermore, they assert that previous mutagenesis studies found synonymous mutations to be far less harmful than nonsynonymous mutations in humans and yeast.

In this article, we assess the validity of Kruglyak *et al*.’s hypothesis and Dhindsa *et al*.’s claims.

## RESULTS

### Testing Kruglyak *et al*.’s hypothesis

Kruglyak *et al*. hypothesize that our estimates of fitness effects of synonymous and nonsynonymous mutations in yeast largely reflect the fitness effects of CRISPR/Cas9 off-target edits and secondary mutations that occurred in mutant construction [2]. While off-target edits are known in plants and animals, the rate of such edits is low and well-designed experiments can avoid off-target edits even in the human genome [4]. The genome is 250 times smaller and the efficiency of non-homologous end-joining relative to homologous recombination (a predictor of off-target editing [5]) is drastically lower [6] in yeast than in humans, rendering off-target editing negligible in yeast. Indeed, no off-target edits were found by genome sequencing in previous yeast studies [7,8]. In our study, gRNAs were carefully designed using Benchling (www.benchling.com/crispr) to minimize potential off-target editing.

Although secondary mutations are possible, the mutation rate of the wild-type yeast used is only ∼0.002 per genome per generation [9]. None of our 21 genes are among those that increase the mutation rate upon deletion (including transformation) found in a previous genome-wide screening [10]. Furthermore, Kruglyak *et al*.’s belief that only a minority of nonsynonymous mutations are significantly deleterious implies that many potential secondary mutations would have no detectable fitness effects, contrasting their Fig. 1. Note that the reported secondary genetic alterations in the yeast deletion collection resulted from selection for beneficial mutations that suppress the harm of gene deletion [11], so cannot explain the “lower-than-expected” mutant fitness [2].

**Fig. 1.**
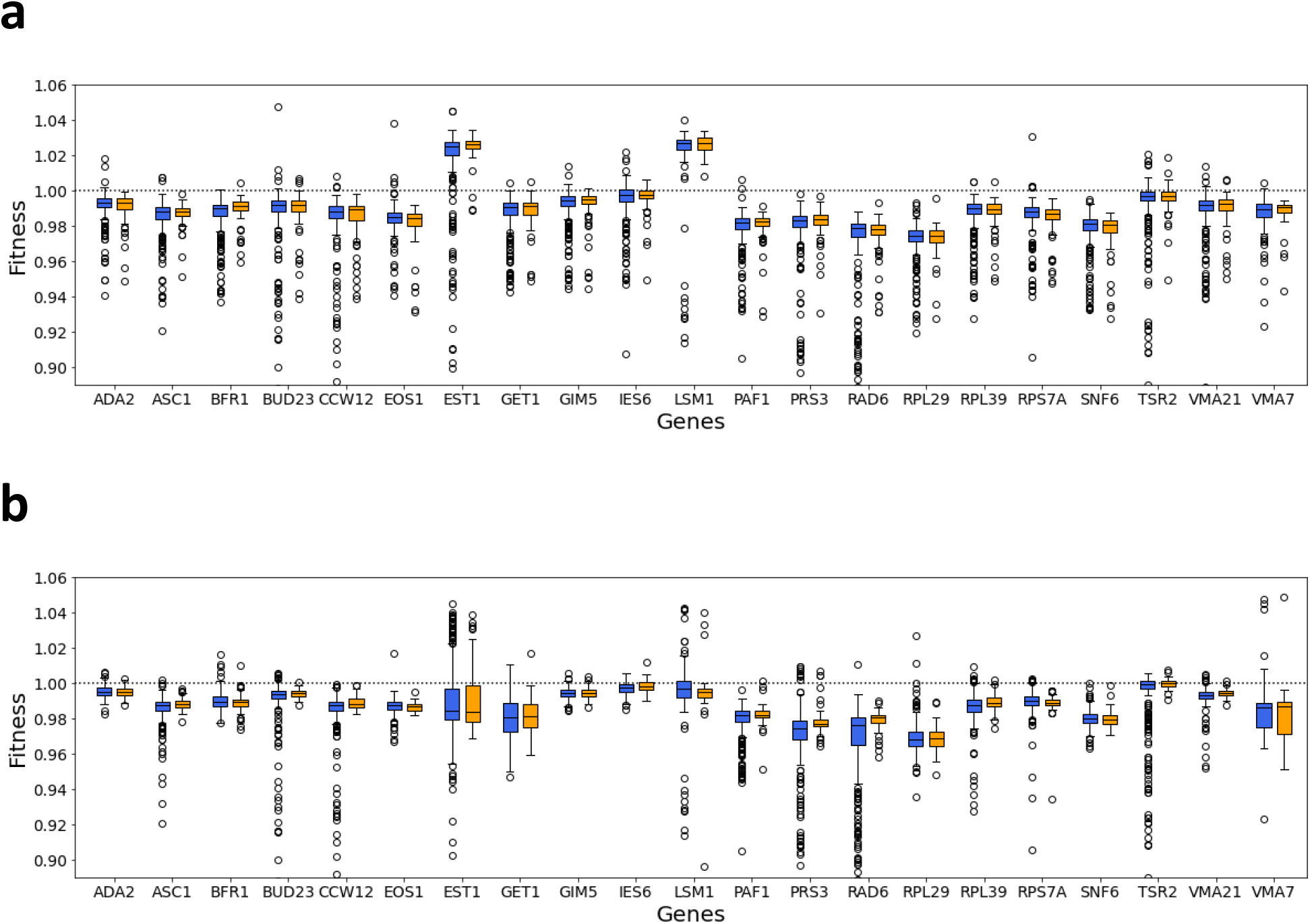
Simulated and experimentally measured fitness of synonymous (yellow) and nonsynonymous (blue) mutants of 21 yeast genes. **a**, Fitness simulated under Kruglyak *et al*.’s hypothesis. To be conservative in testing Kruglyak *et al*.’s hypothesis, we set the simulated fitness values of nonsynonymous mutants that are less fit than all synonymous mutants of the same gene at the values estimated in [1]. **b**, Estimated fitness from [1]. The lower and upper edges of a box represent the first and third quartiles, respectively, the horizontal line inside the box indicates the median, the whiskers extend to the most extreme values inside inner fences (median ± 1.5× interquartile range) and the dots show outliers. Panel b is redrawn using the data in [1]. See Methods for details of the simulation.

The above considerations suggest little if any effect on our mutant fitness measurement from potential off-target edits and secondary mutations. Consistently, all three colonies of the wild-type control constructed through two rounds of CRISPR/Cas9 editing are equally fit as the original wild type (Extended Fig. 1c in [1]).

Kruglyak *et al*. are correct that each mutant is made up of multiple (∼25 on average) independently edited cells in our mutant pool, and the fitness of the mutant measured via *en masse* competition is the average fitness of these cells. They hypothesized the existence of an ascertainment bias in picking relatively healthy colonies for growth rate measurement, predicting that (1) the fitness of a mutant measured through *en masse* competition should be lower than that measured by the growth of a hand-picked colony and (2) the two fitness estimates should not be well correlated due to the influences of different off-target edits and secondary mutations. These predictions are unsupported by our data, because the two fitness estimates show a strong correlation (*r* = 0.90, *P* = 1.3×10^−9^; see Fig. 1d in [1]) with the slope of the linear regression not significantly different from 1 (*P* > 0.2). Kruglyak *et al*. commented that several mutations appear beneficial based on the growths of hand-picked colonies but deleterious based on the *en masse* competition (Fig. 3 in [2]). In fact, only three mutations fit this criterion and the growth rate-based fitness effect is not significantly positive for any of them. Note that the two methods of fitness estimation are equally sensitive for the mutants in Fig. 1d of [1] (average standard error *SE* = 0.00374 and 0.00377 for the X- and Y-axes of fitness estimates, respectively).

Kruglyak *et al*.’s hypothesis makes several predictions on the distribution of fitness effects (DFE) of mutations. First, if synonymous mutations have no genuine fitness effects, the dispersion of the DFE of the synonymous mutations of a gene is caused entirely by off-target edits and secondary mutations in the second round of CRISPR/Cas9 editing that used the same gRNA for all genes, hence is expected to be equal for the 21 genes studied. However, the dispersion (e.g., the interquartile range) varies greatly across the 21 genes (Fig. 2c in [1]). Is this variation explainable by chance given that the number of synonymous mutants of a gene is limited? Using the observed fitness effects of synonymous mutations, we performed a computer simulation to create the DFE of synonymous mutations of each gene under Kruglyak *et al*.’s hypothesis (see Methods). These simulated DFEs (**Fig. 1a**) are highly similar across genes and distinct from the observed DFEs (**Fig. 1b**) in the interquartile range, contradicting the prediction of Kruglyak *et al*.’s hypothesis [2].

**Fig. 2.**
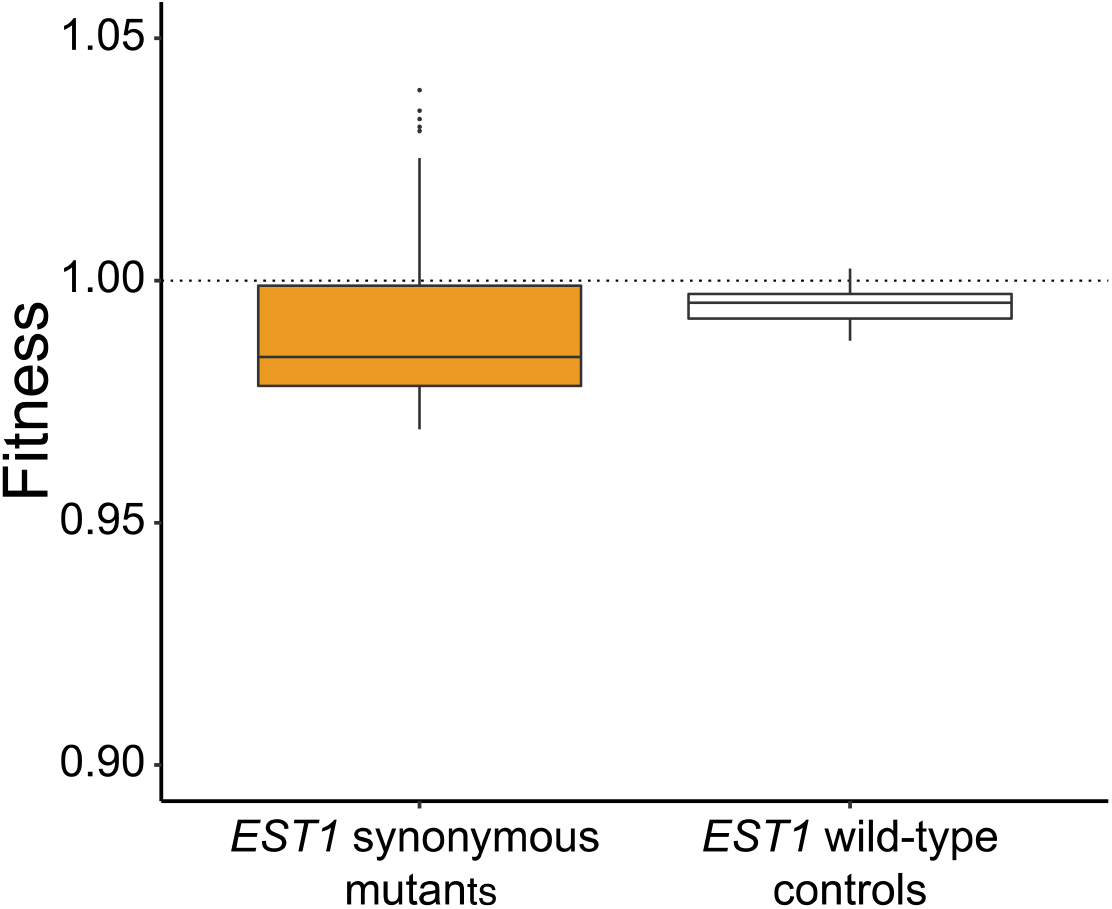
Fitness distributions of 65 synonymous mutants (yellow) and 65 wild-type controls (white) of *EST1*, respectively. The lower and upper edges of a box represent the first and third quartiles, respectively, the horizontal line inside the box indicates the median, the whiskers extend to the most extreme values inside inner fences (median ± 1.5× interquartile range) and the dots show outliers. The wild-type control data are generated from *N*(0.995, 0.003^2^), as explained in the main text.

Second, Kruglyak *et al*. believed that only a minority of nonsynonymous mutants, especially those showing fitness lower than that of almost all synonymous mutants, are truly deleterious, while the rest are neutral. We thus simulated DFEs of nonsynonymous mutations following their hypothesis (see Methods). Again, the interquartile range of the simulated DFE of nonsynonymous mutations is highly similar across the 21 genes (**Fig. 1a**), contrasting the large variation observed (**Fig. 1b**). Furthermore, the interquartile range of the DFE of nonsynonymous mutations is always close to that of synonymous mutations of the same gene in the simulated data (**Fig. 1a**), unlike the pattern in the actual data (**Fig. 1b**).

Third, Kruglyak *et al*. believed that nonsense mutants of the same gene should all have the same fitness, so considered the observed relatively large fitness variation evidence for the hypothesized effects of off-target edits and secondary mutations. Their hypothesis predicts that (1) the DFE of nonsense mutations has the same dispersion across the 21 genes and (2) the DFEs of synonymous and nonsense mutations have equal dispersions, both squarely contradicted by the observation (Fig. 4 in [2]). Two factors explain the relatively large dispersion of the DFE of nonsense mutations. First, because of the relatively low fitness of nonsense mutants, numbers of sequencing reads of these mutants after the competition tend to be low, increasing the sampling errors of the fitness estimates—the average *SE* of fitness is 0.006 (0.012 when fitness < 0.90) for nonsense mutants as opposed to 0.002 for synonymous mutants. Second, transposon mutagenesis suggested that nonsense mutations at different positions of a gene can have drastically different fitness effects and that the relationship between the position and effect can be complex [12].

We also collected new data to directly test Kruglyak *et al*.’s hypothesis. Specifically, from the *ΔEST1* strain used for constructing all coding mutants of *EST1*, we reconstructed wild-type controls and randomly picked 27 colonies. According to Kruglyak *et al*., the average fitness of the 27 colonies is an ideal neutral benchmark against which all *EST1* mutants should be compared, because, as mentioned, each mutant in our *en masse* competition was made up of ∼25 independently edited cells. We measured the growth rate of each of the 27 colonies once and found their average fitness relative to the wild-type control used in the *en masse* competition [1] to be 0.995 (*SE* = 0.003). The fitness of the *EST1* wild-type control is not significantly different from 1 (*P* = 0.16, *t*-test). Importantly, our data reject the prediction from Kruglyak *et al*.’s hypothesis that the variance of the estimated fitness of synonymous mutants of *EST1* equals that of wild-type controls (*F* = 0.018^2^/0.003^2^ = 36, *P* < 10^−13^). This is also visually clear when the 65 synonymous mutants of *EST1* investigated in our study [1] are compared with an equal number of wild-type controls (generated from a normal distribution with mean = 0.995 and variance = 0.003^2^) (**Fig. 2**). Furthermore, 78.5% of *EST1* synonymous mutations are significantly non-neutral when compared with the new wild-type control (nominal *P* < 0.05, *t*-test), similar to that (81.5%) in [1].

Kruglyak *et al*. asserted that the deleterious mutations we reported should have been selectively purged and have no chance to be observed in other yeasts. This inference is based on the unrealistic assumption that (1) the natural environment of yeast is identical to the lab environment and constant and (2) epistasis is minimal. In fact, deleterious mutations in one species are sometimes fixed in another species [13-16]. Our finding that (synonymous and nonsynonymous) mutations unobserved in other yeasts are on average more detrimental in *S. cerevisiae* than those observed [1] provides unequivocal evidence that our measured fitness effects are biological.

Kruglyak *et al*. commented that the mean relative expression level (*REL*) of all mutants of a gene is not correlated with the mean fitness of the mutants of the gene in our data (their Fig. 2). As is clear from Fig. 3c in [1], the mean *REL* is irrelevant because the correlation between *REL* and (rescaled) fitness differs for *REL* <1 and *REL* >1. Furthermore, because gene deletion (i.e., a 100% expression reduction) has different fitness effects for different genes, using rescaled fitness is essential in analysis across genes. It is worth emphasizing that the correlation between *REL* and rescaled fitness across all mutants with *REL* < 1 is moderate (*ρ* = 0.32 and 0.30 for synonymous and nonsynonymous mutants, respectively [1]). This fact, coupled with the observation that the *REL* distribution for a gene can be skewed depending on the gene expression level (Extended Data Figs. 4i, 5f, 5h in [1]), means that the correlation between *REL* (when *REL* < 1) and fitness may or may not be significant for individual genes. Note that even when the above correlation is not significant, it does not mean that the fitness estimates are wrong because coding mutations can affect fitness via multiple mechanisms [1].

Together, the above analyses reject Kruglyak *et al*.’s hypothesis as an explanation of our data. Kruglyak *et al*. commented that our results are inconsistent with a large body of literature, without realizing that the prior results were almost exclusively based on indirect inferences of fitness effects that depended on many simplifying assumptions that may or may not hold and that the analysis of prior mutagenesis studies by Dhindsa *et al*. is seriously flawed (see below).

### Evaluating Dhindsa *et al*.’s analyses and interpretations

In their Fig. 1, Dhindsa *et al*. [3] presented the cumulative probability distributions of minor allele frequencies (MAFs) of several types of mutations observed from 454,668 human exomes, noting that MAFs are overall higher for synonymous than nonsynonymous mutations. They interpret this disparity as weaker purifying selection on synonymous than nonsynonymous mutations. We do not disagree with this interpretation but note the following important caveats. First, the infinite site model is violated when the sample size is so large [17]. Consequently, the mutation rate is a primary determinant of the MAF distribution [17]. Because transitions (especially CpG → TpG) occur more frequently than transversions and tend to be synonymous [17], synonymous mutations look less constrained in the MAF distribution than they actually are. In other words, the true difference in *s* between synonymous and nonsynonymous mutations is smaller than what their Fig. 1 implies. Second, the MAF distributions largely reflect the coefficients of selection against deleterious mutations in the heterozygous state, while the mutational fitness effects were measured in haploid yeast in our experiment [1]. If dominance differs between synonymous and nonsynonymous mutations (e.g., synonymous mutations are completely recessive but nonsynonymous mutations are partially recessive), the MAF distributions could look different from those expected from our haploid experimental results. Third, selections inferred from MAF distributions are not fundamentally different from those inferred from nonsynonymous to synonymous substitution rate ratios (*d*_N_/*d*_S_). We discussed extensively scenarios under which observations of *d*_N_/*d*_S_ <<1 can be reconciled with our experimental estimates of fitness effects [1], and the same applies here. Fourth, while we emphasized the strong deleterious effects of synonymous mutations, our data showed that nonsynonymous mutations are overall significantly more deleterious than synonymous mutations in each of the four environments examined [1]. Hence, the human MAF distributions may not be inconsistent with our experimental results, especially when all of the above factors are considered. Finally, we note that “neutral synonymous” (grey line) in Fig. 1 of [3] does not mean neutrality in fitness but neutrality in codon stability coefficient, so using this terminology when discussing fitness effects is misleading.

In their Fig. 2, Dhindsa *et al*. [3] reported that, using gene-based collapsing analysis performed on 394,694 UK Biobank participants, they found 2, 32, and 55 genes to be associated with at least one clinical phenotype when only synonymous, nonsynonymous, or nonsense mutations were considered, respectively. However, this comparison is unfair because the authors restricted their analysis to nonsynonymous mutations predicted to be pathogenic (REVEL score [18] ≥ 0.25) while no similar restriction was set for synonymous mutations. Furthermore, the very low numbers of genes identified to be phenotypically associated indicate that this analysis is severely underpowered; it underestimates the phenotypic contributions of all types of mutations so cannot inform the general clinical relevance of synonymous mutations. It is entirely possible that the relative importance of synonymous and nonsynonymous mutations to disease shows a pattern substantially different from that in their Fig. 2 when (1) a fair comparison is made, (2) the statistical power is greatly increased, and (3) a much larger proportion of variants are examined (Fig. 2 analyzed only extremely rare variants). Indeed, using single-nucleotide resolution quantitative trait locus mapping, She and Jarosz mapped many yeast growth traits to synonymous variants and found the effect size comparable between synonymous and nonsynonymous causal variants [19].

It is worth noting that the relationship between the impact of a mutation on disease and that on fitness is complex and nonlinear. For example, a life-threatening disease may have a minimal fitness effect if it occurs after the reproductive age. By contrast, a mutation that lowers the fertility (and hence fitness) by 5% probably will not even be considered disease-causing if it has no other phenotypic effect. It is thus also possible that synonymous and nonsynonymous mutations are significantly different in clinical relevance given their significant difference in fitness effect, especially that mutations with *s* < -0.05 are almost exclusively nonsynonymous in our data [1].

In Extended Data Fig. 1, Dhindsa *et al*. [3] presented a comparison between synonymous and nonsynonymous mutations using multiplexed assays of variant effect (MAVEs) for seven human genes (instead of eight mentioned in their Methods) and two yeast genes. The lower median “score” of nonsynonymous than synonymous mutants and the proximity of the median score of synonymous mutants to the “no effect” line appear to suggest that nonsynonymous mutations are substantially more deleterious than synonymous mutations and that synonymous mutations are largely neutral. A closer examination of the data (MaveDB [20]) behind this comparison raises many red flags. Three problems are shared between the human and yeast data analyzed. First, many nonsynonymous mutants each contained multiple nonsynonymous mutations while each synonymous mutant appears to contain only one synonymous mutation [21-27], making their comparison unfair. Second, mutant genes were not in their native chromosomal locations but were placed on plasmids [21-25,27]; plasmid copy number variations affect mutant protein abundance and interfere with the quantification of the mutational effect. Third, the measurement error was large in at least some datasets, reducing the likelihood of detecting mutational effects. For example, the average standard error of the mutant score in the human *TPK1* dataset (urn:mavedb:00000001-d-1) was 0.09 (in the scale from 0 to 1). If the score is equivalent to fitness, a mutant must have a fitness lower than 0.82 for the mutation to be called significantly deleterious. By this standard, no synonymous mutations (but several nonsynonymous mutations) in our data would be significantly deleterious.

The human data analyzed by Dhindsa *et al*. [3] additionally suffered from the following problems. First, in four (*UBE2I, SUMO1, CALM1*, and *TPK1*) of the seven genes studied, mutants were examined in yeast by a complementation assay [21]; to what extent the mutational effects measured in yeast are relevant to humans (or even human cells) is unknown. Furthermore, the scores were normalized to a scale from 0 to 1, where 0 corresponded to the median score of nonsense mutants while 1 had different meanings in different datasets. In one dataset (00000001-a-4), 1 corresponded to the wild type. However, in all other datasets (00000001-a-3, 00000001-b-2, 00000001-c-1, and 00000001-d-1), 1 corresponded to the median score of synonymous mutants, artificially making synonymous mutations “neutral”. Second, for the fifth and sixth human genes studied (*PTEN* and *TMPT*), the score reflected the protein expression level rather than fitness [26]. Finally, for the seventh human gene (*CYP2C9*), the score was again not fitness but enzyme activity measured in yeast cells [27] (in dataset 00000095-a-1) or protein abundance measured in a human cell line [27] (in dataset 00000095-b-1).

All yeast data analyzed by Dhindsa *et al*. [3] additionally had the problem of using non-native promoters to drive the expression of mutant genes. For example, in all three datasets of *HSP90* (00000011-a-1, 00000039-a-1, and 00000040-a-1), mutant gene expression was driven by the *GPD* promoter [22,24] or *ADH* promoter [23]. Similarly, in the *UBI4* dataset (00000037-a-1), mutant gene expression was driven by the *GPD* promoter [25]. The biological relevance of the fitness effects measured under these non-native promoters is unknown.

It is therefore clear that Extended Data Fig. 1 in [3] does not provide valid information on the fitness effects of human or yeast synonymous and nonsynonymous mutations. Additionally, under Kruglyak *et al*.’s hypothesis, secondary mutations would have affected these studies as well.

## DISCUSSION

Our analysis of published and newly collected data demonstrates that Kruglyak *et al*.’s hypothesis cannot explain our estimates of fitness effects of mutations in yeast. Kruglyak *et al*. recommended that the wild-type control be reconstructed along with all mutants in the same large-scale CRISPR/Cas9 editing experiment. We had considered this option in our previous study, but because (1) the large-scale editing experiment cannot guarantee the generation of all designed genotypes (e.g., we were able to estimate the fitness of only ∼90% of designed mutants), (2) the reconstructed wild type is required in our study, and (3) off-target editing is negligible and secondary mutations are rare in yeast, we chose to reconstruct the wild type separately to ensure its presence in the *en masse* competition. This said, the risk of not generating the reconstructed wild type in the large-scale editing experiment can be minimized by increasing the concentration of the wild-type template DNA in the experiment. Hence, we agree with Kruglyak *et al*. on their recommendation when off-target editing and secondary mutations are of concern.

We found that, when examined critically, Dhindsa *et al*.’s results are not inconsistent with our estimates of fitness effects of yeast synonymous and nonsynonymous mutations. Specifically, their comparisons between synonymous and nonsynonymous mutations in prior mutagenesis studies and in contributions to disease are unfair and human allele frequency distributions can be compatible with our fitness estimates when multiple complicating factors are considered.

It is worth mentioning that recent years have seen an increasing number of reports of fitness effects of synonymous mutations from both case studies [28-32] and systematic analyses [19,33-35]. Notably, Lind *et al*. found similarly large fitness effects of synonymous and nonsynonymous mutations in two ribosomal protein genes of the bacterium *Salmonella typhimurium* [34]. Sane *et al*. found synonymous and nonsynonymous mutations observed from *Escherichia coli* mutation accumulation experiments to have comparable fitness effects [35]. Sharon *et al*. reported similar fractions of synonymous and nonsynonymous differences between two yeast strains to have detectable fitness effects [33]. She and Jarosz mapped many yeast growth traits to synonymous variants and discovered their substantial causal growth effects [19]. These findings, along with ours [1], suggest that many synonymous mutations are strongly non-neutral. More studies are certainly needed to examine the generality of these findings across the tree of life and explore its potentially very broad implications. A number of synonymous mutations have already been reported to cause disease [36], and a systematic survey will only find more cases.

## METHODS

### Simulation of DFEs under Kruglyak *et al*.’s hypothesis

We simulated DFEs of synonymous and nonsynonymous mutations following Kruglyak *et al*.’s hypothesis. Specifically, the simulation assumed that all synonymous mutations’ fitness effects are due to errors (i.e., off-target edits and secondary mutations). We separately inferred the fitness effects of errors from the two rounds of CRISPR/Cas9 editing (error_I and error_II, respectively).

Error_I is the same for all synonymous mutants of a gene, because they were all made from the same gene-deletion strain created by the first round of CRISPR/Cas9 editing. For a gene, we chose the 95th percentile of its synonymous DFE as the value of its error_I, under the assumption that 5% of synonymous mutants are made fitter while 95% are made less fit by errors in the second round of CRISPR/Cas9. The simulation outcome (in terms of the dispersion of the DFE) is insensitive to the above percentile choice.

Because the second round of CRISPR/Cas9 is the same for all genes, we expect error_II to follow the same distribution for all mutants of all genes. Under the assumption that each synonymous mutation’s fitness effect equals error_I + error_II, we estimated error_II of each synonymous mutation. From all synonymous mutations of all genes, we obtained a probability distribution of error_II.

We then simulated the fitness effect of each synonymous mutation by adding its gene-specific error_I and a random variable sampled from the probability distribution of error_II. This was done for every synonymous mutation in our data to create the simulated DFEs of synonymous mutations of the 21 genes.

Kruglyak *et al*. contended that only nonsynonymous mutations that are more deleterious than almost all synonymous mutations are genuinely deleterious. Let *A*_*i*_ be the minimal fitness value of all synonymous mutants of gene *i*. For a nonsynonymous mutant of gene *i* with fitness ≥ *A*_*i*_, we simulated its fitness the same way we simulated the fitness of a synonymous mutant of gene *i*; otherwise, the simulated fitness equals its observed fitness. This was done for every nonsynonymous mutant in our data to create the simulated DFEs of nonsynonymous mutations of the 21 genes.

### Fitness distribution of *EST1* wild-type controls

The experiments generally followed [1]. We amplified the wild-type *EST1* gene from the genome of the haploid strain BY4742 by polymerase chain reaction (PCR) using the high-fidelity Q5 polymerase (NEB) and inserted it into the *ΔEST1* cells used in [1] by CRISPR/Cas9 editing. Twenty-seven colonies were randomly picked and the insert was confirmed by the PCR product length and Sanger sequencing for each of them. The cells were then counter-selected on 5-FOA plates to remove the CRISPR/Cas9 plasmids. We measured the maximum growth rate of each of the 27 strains by monoculture [1], with one replicate per strain. We also similarly measured the growth rate of the wild-type control used in [1] with 31 replicates. The fitness of each of the 27 strains relative to the wild-type control used in [1] was calculated as in [1]. The average fitness of the 27 strains as well as its *SE* were then computed. We compared the average fitness of the 27 strains with that of the 31 replicates of the wild-type control by a *t*-test. Because we measured the growth rate of each of the 27 strains only once, the *SE* is likely larger than its expected value in *en masse* competitions, making our conclusion that the *SE* (which equals the standard deviation of fitness among multiple wild-type controls) is smaller than the standard deviation of the fitness distribution of synonymous mutants of *EST1* conservative.

## ACKNOWLEDGMENTS

We thank Haiqing Xu for discussion. This work was supported by the U.S. National Institutes of Health research grant R35GM139484 to J.Z.

